# Factors affecting the individual probability of infection with a prevalent pathogen (*Mycoplasma*) and the effect on Griffon vultures’ movement behavior

**DOI:** 10.1101/2024.08.01.606137

**Authors:** Nili Anglister, Miranda May Crafton, On Avraham, Marta Acácio, Gideon Vaadia, Ohad Hatzofe, Yigal Miller, Inna Mikula, Noa Pinter-Wollman, Inna Lysnyansky, Orr Spiegel

**Affiliations:** School of Zoology, Faculty of Life Sciences, Tel Aviv University, Tel Aviv, Israel; Koret School of Veterinary Medicine, the Robert H. Smith Faculty of Agriculture, Food and Environment, the Hebrew University of Jerusalem, Israel; Science Division, Israeli Nature and Parks Authority, Jerusalem, Israel; Division of Avian Diseases, Kimron Veterinary Institute, POB 12, Bet Dagan 50250, Israel; Department of Ecology and Evolutionary Biology, University of California Los Angeles, Los Angeles, California, USA

## Abstract

Mycoplasmas are known as commensals and pathogenic bacteria of various raptor species causing clinical or subclinical infections. However, little is known about the prevalence of mycoplasma in captive and wild raptors and its significance to their health. In Israel, the Griffon vulture (*Gyps fulvus*; hereafter Griffons) is considered critically endangered, and its intensive management program includes population monitoring and restocking (captive-born or imported rehabilitated wild Spanish Griffons). Here we survey the prevalence of *Mycoplasma* species in both the wild and captive populations. During 2019-2020, we collected 244 tracheal swab samples from 167 unique individuals. We used PCR analysis to identify *Mycoplasma* species. First, we identified nine spp., including species not yet described in Israel or for Griffon vultures. Second, imported Griffons showed a higher prevalence and a different diversity of species in comparison to the local ones, suggesting that at least one *Mycoplasma* species (Sp 18b) was introduced into the native population. Third, juvenile Griffons had a higher prevalence, different species composition, and stronger reduction in movement compared to adults, confirming the susceptibility of this group to mycoplasma infections. GPS-tracking of 60 free-ranging individuals showed that even in the absence of apparent clinical signs, Griffons infected with mycoplasma, and especially sub-adults flew less (shorter distances and periods). These findings underscore the importance of considering potential pathogen introductions in population reinforcement and reintroduction initiatives, providing valuable insights for similar conservation programs globally. Further, they demonstrate the potential of long-term tracking for detecting subclinical effects that are unnoticeable in clinical examination.

## 1. Introduction

Disease risk assessment is extremely important in wildlife management in general and reintroduction programs in particular. Although sometimes overlooked, diseases may result in expensive failures of management efforts, and worse still, the introduction of destructive pathogens into naïve resident wildlife populations (Aguirre 2002; Kock *et al*. 2010). Stress endured in captivity, the close proximity of captive-raised individuals, as well as genetic effects (inbreeding or drift), may increase pathogen and parasite prevalence in reintroduction candidates (Mason 2010; Ewen *et al*. 2012). The transfer of individuals to a new location can cause transmission of novel pathogens into the environment and cause spillovers into other wildlife species, as well as domestic animals and humans (Kock *et al*. 2010; Ballou 2013). Some notable examples include the spread of *Plasmodium* spp. to extant wild turkeys (*Meleagris gallopavo*) (Castle & Christensen 1990), the spread of *Mycoplasma gallisepticum* to house finches (*Haemorhous mexicanus*) (Fischer *et al*. 1997; Dhondt *et al*. 1998; Hochachka *et al*. 2013); and of foot and mouth disease virus spilling over from translocated Cape buffaloes (*Syncerus caffer*) to cattle penned near the relocation site (Hedger & Condy 1985). Conversely, exposure of translocated animals to a wide range of new pathogens they have not developed immunity to has led to several recorded morbidity and mortality events, impeding the program success. Notable examples include: reintroduced Arabian oryx (*Oryx leucoryx*) bred in captivity in the United States died of botulism when they were released into the desert of Oman, where this disease is enzootic in sheep and goats (Stanley-Price 1989); near failure of the reintroduction of the black-footed ferret (*Mustela nigripes*) in the United States, due to their contracting canine distemper (Williams *et al*. 1988). These cases highlight the importance of an evaluation of pathogen prevalence in translocation programs to ensure the success and sustainability of wildlife management efforts.

Pathogens affect their hosts in several ways, potentially influencing host fitness, habitat preference, social interactions and behavior (Ezenwa *et al*. 2016; Binning *et al*. 2017; Spiegel *et al*. 2022) sometimes to benefit the pathogen’s survival (Poulin & Maure 2015; Godfrey & Poulin, R. 2022). Pathogens often reduce an individual’s locomotion through morbidity (lethargy) or altering its physiology and physical condition (Goodman & Johnson 2011; Binning *et al*. 2017). For example, Pacific chorus frogs infected by trematodes suffer from severe limb malformations which directly decreases swimming and jumping distance (Goodman & Johnson 2011). Pathogens may also cause a deterioration in body condition, reducing energy reserves and thus locomotion capacity. This has been demonstrated in multiple studies, such as mange-infected wolves, vultures with avian influenza or tick-infected lizards (Main & Bull 2000; Cross *et al*. 2016; Duriez *et al*. 2023). In addition, pathogens may affect host movements by influencing their ability to acquire or process information on their environment. One example of such host manipulation is *Toxoplasma gondii* infection in rats (*Rattus norvegicus*), which increases activity that exposes them to cats (the main reservoir of this parasite), due to an alteration in the rat’s perception of predation risk (Vyas *et al*. 2007; Hari Dass & Vyas 2014). By using advanced telemetry technology, we can account for the role of disease on heterogeneous individual behavior in threatened species (e.g. their movement and social contacts) and on their risk of being exposed to different threats (Kays *et al*. 2015; Dougherty *et al*. 2018; Spiegel *et al*. 2022; Duriez *et al*. 2023).

Not all microorganisms will affect the hosts fitness and behavior. Microorganisms are usually separated into commensals, which typically coexist with their host without causing harm (Dobrindt *et al*. 2013), mutualists, e.g. the gut flora of performs essential functions by processing undigested food to the benefit of the host and limiting colonization of the gastrointestinal (GI) tract by pathogens (Gilmore & Ferretti 2003); and pathogens, which can cause damage in a host, often leading to disease (Pirofski & Casadevall 2012; Hornef 2015). However, the distinction between the two is not always clear-cut, as some commensal microorganisms can become pathogenic under certain conditions. For example, gut bacteria, which are usually beneficial, can become pathogenic when they acquire new traits, such as overt virulence and antibiotic resistance (Gilmore & Ferretti 2003; Conway & Cohen 2015). Therefore, the definition of a pathogen compared to a mutualist or commensal microorganism is sometimes context dependent, affected also by the microorganism’s behavior and the host’s immune response (Gilmore & Ferretti 2003; Henriques-Normark & Normark 2010; Dobrindt *et al*. 2013; Hornef 2015).

Here we focus on *Mycoplasma* spp. in Griffon vultures (*Gyps fulvus*, hereafter Griffons) in Israel as a model of a commensal/pathogen in an extensively managed locally endangered species. *Mycoplasma* is a genus of bacteria that lack a cell wall around their cell membranes (Razin *et al*. 1998) and is the smallest self-replicating life form. They cause various diseases in humans, domestic animals, and wildlife (Sumithra *et al*. 2013; Dawood *et al*. 2022). Mycoplasmas may have harmful effects in both free-ranging and captive reptiles, mammals and birds (Fischer *et a*l. 1997; Sumithra *et al*. 2013). They can cross between wild and domestic species (Giacometti *et al*. 2002; Arif *et al*. 2007; Ostrowski *et al*. 2011; Highland *et al*. 2018) and can also be zoonotic (Baker *et al*. 1998; Friend 2006; Lierz *et al*. 2008b; Prayson *et al*. 2008). In wild ruminants, the emergence of different *Mycoplasma* species has caused diseases and has harmed populations of many species (Black *et al*. 1988; Verbisck-Bucker *et al*. 2008; Janardhan *et al*. 2010; Besser *et al*. 2012; Malmberg *et al*. 2020).

In reptiles, *Mycoplasma* spp. cause chronic upper respiratory tract disease (URTD) in captive and free-ranging tortoise populations including desert tortoises (*Xerobates agassizii* ) and Gopher tortoises (*Gopherus polyphemus* ), which is jeopardizing these endangered species (Jacobson *et al*. 1991; Brown *et al*. 1994, 2004a; Jacobson *et al*. 2014). The release of tortoises held in captivity from rehabilitation or breeding programs are thought to be the source for this disease in free-ranging populations (Jacobson *et al*. 1991; Brown *et al*. 2002) emphasizing the importance of monitoring mycoplasma in conservation programs and breeding programs and before release of captive individuals into the wild. *Mycoplasma* spp. were also found to cause fatal disease (pneumonia, arthritis and hepatitis) in Nile Crocodiles (*Crocodylus niloticus*) in Zimbabwe (Mohan *et al*. 1995, 1997), and American alligators (*Alligator mississippiensis*) and caimans (*Caiman latirostris*) in Florida (Clippinger *et al*. 2000; Brown *et al*. 2004b).

Specifically in wild birds, many different *Mycoplasma* spp. have been isolated; some are thought to be commensal, but some are known to be pathogenic causing conjunctivitis, respiratory infections, arthritis, embryonic death, skeletal deformations and reduced hatchling sizes (Gerlach 1994; Erdélyi *et al*. 1999; Lierz *et al*. 2000a, 2008a; Sumithra *et al*. 2013). *M. gallisepticum*, which is a known pathogen in poultry spread rapidly and caused a great decline in eastern house finch (*Haemorhous mexicanus*) populations in the US (Fischer *et al*. 1997; Dhondt *et al*. 1998; Hochachka *et al*. 2013). In the Native western populations of finches the epidemic expanded slower than in the eastern introduced range (Dhondt *et al*. 2006), and prevalence was found to be higher in young birds (Altizer *et al*. 2004). Mycoplasma has also have been found to cause disease in raptors (Poveda *et al*. 1990a; Panangala *et al*. 1993; Erdélyi *et al*. 1999; Ruder *et al*. 2009; Sumithra *et al*. 2013), mostly in younger raptors (Poveda *et al*. 1990a; Lierz *et al*. 2000b, 2002, 2008a). However, the significance of many species of *Mycoplasma* to raptor health, is yet to be determined (Lierz *et al*. 2000a, 2008a, 2019; Loria *et al*. 2008; Lecis *et al*. 2016). New *Mycoplasma* species are still being found in vultures and in other raptors (Loria *et al*. 2008; Lecis *et al*. 2010; Suárez-Pérez *et al*. 2012).

In this multidisciplinary analysis we combined the (genetic) identification of *Mycoplasma* species from Griffon’s samples with their GPS-tracking to quantify movement behavior. This approach enabled us to provide unique insights when exploring pathogen-host relationships and to make recommendations for future conservation management of Griffons and other endangered species. Our objectives were to:

A. *Examine the effect of the site of sampling and individual traits such as age, origin (captive vs. free roaming Griffons) on mycoplasma prevalence*. *Hypothesis: (H. A1) Mycoplasma prevalence will be higher in juvenile Griffons (higher prevalence of mycoplasma has been found in juveniles of other species (Poveda et al. 1990a; Lierz et al. 2000b, 2002, 2008a)*. *and (H. A2) will be higher in imported Griffons (due to low immunity caused by the stress of the transfer, and to the close proximity between individuals in transfer and enclosures)*.
B. *Examine differences in the different Mycoplasma species present in Griffons from different origins and of different ages*. *Hypothesis: Mycoplasma spp. can differ in their age-dependent susceptibility and their geographical distribution. Accordingly, we predicted that species will (H. B1) differ among age groups, and (H. B2) between imported Griffons and local Griffons*.
C. *Examine whether there is an association between disease (e.g., mycoplasma infection) and movement in Griffons*. *Hypothesis: (H. C1) According to previous research we predicted that Mycoplasma spp. in raptors will be mostly commensal and thus will not affect movement. An alternative hypothesis is (H. C2) that while clinical signs are not commonly observed in raptors the infection of mycoplasma might cause changes in behavior, such as decreased ability to fly*.

## 2. Methods

### 2.1. Study Species: Griffon vultures, an endangered species in Israel

The global decline of vulture populations is a pressing concern, with 12 out of the 23-vulture species classified as endangered or critically endangered, and an additional four considered near threatened, primarily due to poisoning (Ogada *et al*. 2012; Buechley & Şekercioğlu 2016). In Israel, the breeding populations of three out of five vulture species have become locally extinct, including *Torgos tracheliotus negevensis*, *Aegypius monachus*, and *Gypaetus barbatus*. Furthermore, the Egyptian vulture (*Neophron percnopterus*) and the Griffon vultures are locally classified as critically endangered (Mayrose *et al*. 2017; IUCN 2022). The Griffon population in Israel has seen a rapid decline, dropping from thousands a century ago to just over 500 individuals a couple of decades ago and currently numbering less than 200 individuals (Mayrose *et al*. 2017; IUCN 2022).

Like most other vulture species, Griffons are large obligate scavengers (2.4-2.7 m wing span, 6- 10 kg mass), that scan large areas through soaring flight (Ruxton & Houston 2004). Once food is detected, individuals aggregate from considerable distances, attracted by the sight of other vultures gliding to a carcass (Jackson *et al*. 2008). Multiple individuals land and feed simultaneously in a scrambling feast, facilitating disease spread. Griffons roost and nest in communal roosts, which can serve as information centers for locating resources (Harel *et al*. 2017), but may also contribute to disease spread among individuals. Roosts are also breeding colonies where pairs will incubate their single egg for 55 days and rear the chick for at least 100 days until fledging. The chick will take 4-6 years to mature and start breeding. This long and slow recruitment rate (‘K life history strategy’) leads to high population-level sensitivity to adult mortality, also because the loss of a parent causes abandonment of nest and death of the offspring, making the conservation of this species challenging (Sarrazin *et al*. 1996; Xirouchakis 2010).

To address the alarming decline in Griffon vulture populations, the Israeli Nature and Parks Authority (INPA) has instituted a comprehensive management program. This initiative encompasses various conservation strategies, including the operation of supplementary feeding stations to provide uncontaminated food, systematic monitoring of breeding populations through annual surveys and observations, and restocking efforts utilizing vultures bred in Israeli captivity and those imported from Spain (Alon & Hatzofe 2016; Mayrose *et al*. 2017). Successful wildlife management programs in Western Europe, particularly in Spain and France, underscore the efficacy of proactive conservation measures (IUCN 2022).

### 2.2. Experimental design

During the non-breeding season (September-November), the INPA performs routine captures of free-roaming Griffons in two different sites: **Sde-Boker** (**Negev** desert) and **Hai-Bar Carmel** (**Carmel**). To determine prevalence of mycoplasmas in the captive population we also sampled individuals from breeding program facilities and acclimation cages (captive Griffons) at the **Hai Bar Carmel** (**Carmel**), **Jerusalem Biblical Zoo** (**Jerusalem**), and **Gamla** nature reserve (**Golan** Heights), details in **Fig. 1**. Handling times were kept to the bare minimum to avoid excess stress. All Griffons were clinically examined by a veterinarian or veterinary assistant and treated or hospitalized if they showed signs of illness/ debilitation. Due to the Corona virus outbreak, no Griffons have been imported since 2020, so we focused our sampling on free-roaming Griffons in the Negev in 2021-2022.

**Figure 1.**
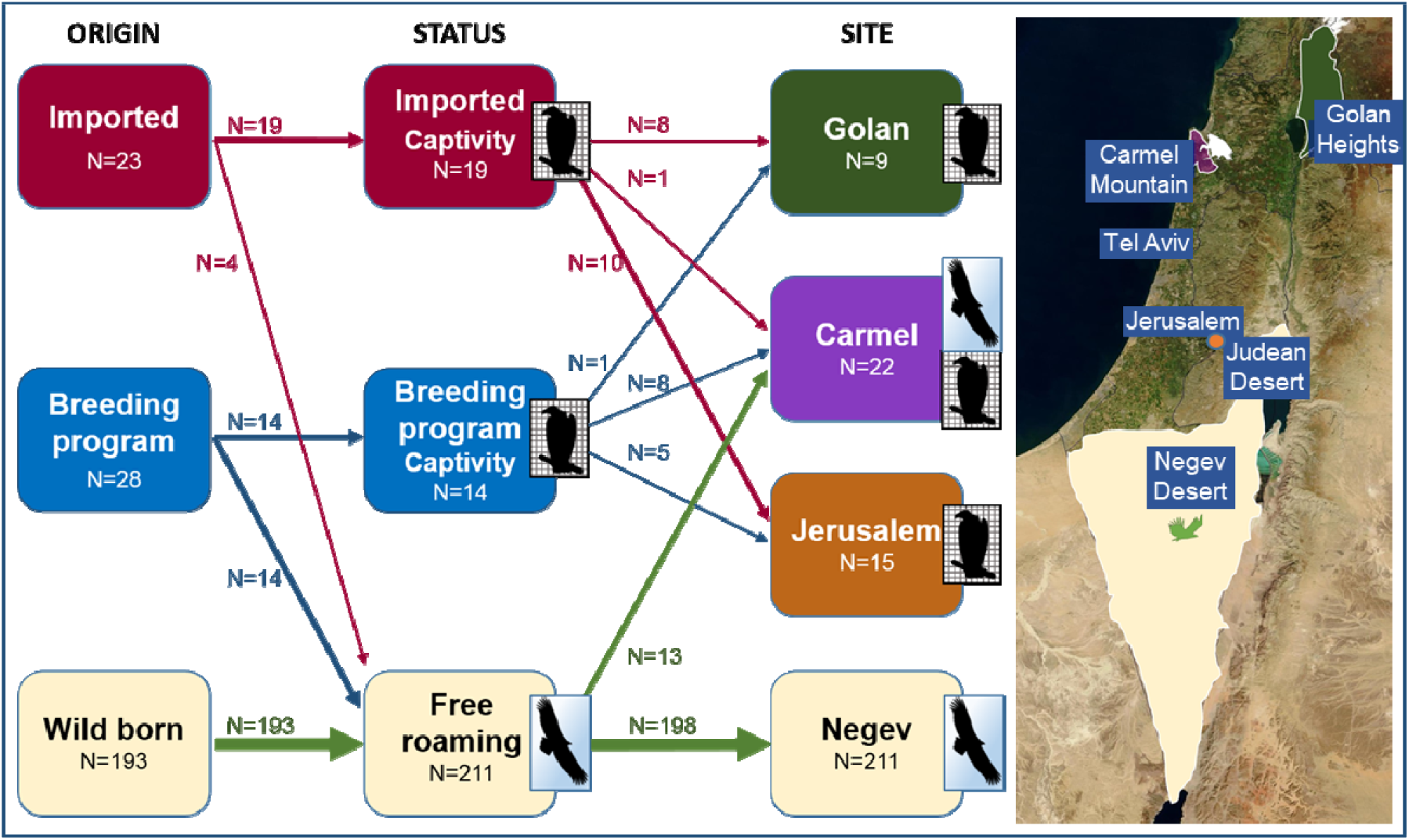
Description of sample design. The left rectangles represent the origins of the sampled Griffons. Red- depicts Griffons imported from Catalonia, Spain (sampled in 2019-2020), blue- breeding program Griffons (sampled in 2019-2020), and yellow wild-born Griffons. The middle rectangles represent the status of the Griffons whether they are free-roaming (yellow) or captive (imported in red and breeding-program in blue) during sampling. The right rectangles represent the site of sampling for the captive Griffons (brown- Jerusalem, green- Golan, purple- Carmel) and the free-roaming (most in the Negev, a few in the Carmel). In the Carmel most of the free-roaming Griffons were from imported origin (from Catalonia, Spain) or breeding program born origin. In all cases, the number of samples for origin, status and site are depicted. Arrow width indicates sample size.

### 2.2. Sampling & Data collection

Samples were taken with a sterile swab from the mucosa of the choana of the Griffons. The samples were kept in -20[C until DNA extraction. Between 2019-2022, we collected and analyzed a total of 244 choanal samples from 167 unique individuals (51 were sampled twice on consecutive years, 12 were sampled three times and one was sampled in all four years), we only analyzed one sample per individual per year for mycoplasma. Swabs were also sent to the Div. Avian Diseases, Kimron Veterinary Inst. (**KVI**), Bet Dagan for identification of New Castle disease virus, West Nile virus, Avian influenza and Chlamydia, in order to assess the vultures’ health.

The following data were recorded for each sample:

1. The identity of the Griffon: according to wing tags, metal rings or color rings
2. The date of sample
3. Sexing and Age: Sex was determined by DNA testing from blood or feathers (Karniely Vet Lab Ltd.). Age was determined from the molting stage or known time of hatching (captive), and grouped for analysis as: *i*) Juveniles- hatching year, *ii*) Sub adults – 1- 4 years post-hatching; *iii*) adults – more than 4 years post-hatching (Zuberogoitia *et al*. 2013).
4. The site of sample: Negev, Carmel, Golan or the Jerusalem zoo.
5. The origin (place of birth, irrespective of sampling location) of the Griffons was defined as: where the Griffon originated from i) Wild born: born in the wild in Israel; ii) Breeding program: born in the breeding program in Israel (i.e., in above-mentioned facilities); iii) Imported: imported from Catalonia to supplement the breeding program and the free-roaming Griffon population.
6. Status of Griffon sampled: where the Griffon was sampled free-roaming or captivity: divided into imported or breeding-program Griffons (**Fig. 1**). Griffons originating from the breeding program or imported to Spain may have been released to the wild (and are free-roaming) or remain in captivity in the breeding program

### 2.3. Genomic DNA extraction & replication for Mycoplasma spp

DNA extraction and replication were conducted in the Div. Avian Diseases, Kimron Veterinary Inst. (**KVI**), Bet Dagan. DNA for template was prepared directly from individual choanal swab agitated vigorously in 1 ml of PBS (Sigma, Rehovot, Israel). Genomic DNA was extracted from 400 µl of PBS solution using the Maxwell DNA Isolation Kit for Cell/Tissue and the Maxwell® 16 apparatus (Promega) according to the manufacturer’s instructions. The identification of mycoplasma species was based on the partial sequence of the 16S rRNA gene (around 1000 bp in length). Extracted DNA was amplified using GPF (5’ GCT GGC TGT GTG CCT AAT ACA 3’ (Lierz *et al*. 2007) and MGSO (5’ TGC ACC ATC TGT CAC TCT GTT AAC CTC 3’ (Van Kuppeveld *et al*. 1992) primers. PCRs were carried out in 25 µl and contained 0.25 µl of Phire Hot Start II DNA Polymerase (Thermo Fisher Scientific, Waltham, MA, USA), x5 Phire reaction buffer, 0.5 µl of 10 mM dNTPs, 0.4 µM each primer and 5 µl DNA. PCR amplifications were carried out in a C1000 Touch™ Thermal Cycler (Bio-Rad, Hercules, CA, USA).

Amplification was conducted as previously described (Lierz *et al*. 2007) with a slight modification as follows: 94 °C for 3 min, followed by 35 cycles of denaturation for 30 sec at 94°C, annealing for 30 sec at 66°C and synthesis for 1 min at 72°C, and then a final extension for 5 min at 72°C. DNA of *M. falconis* and nuclease free water (Sigma, Rehovot, Israel) were used as positive and negative controls, respectively. The amplified PCR products (5 μl) were then separated in a 1% agarose gel and visualized by ethidium bromide staining and ultraviolet transillumination. A biomarker bp-100 Bio-Rad, (Hercules, CA, USA) was used to calculate the size of DNA fragments. Positive PCR samples were then purified using the MEGAquick-spinTM-spin PCR & Agarose Gel DNA Extraction System (iNtRON Biotechnology) and subjected to the sequencing (Hylab Ltd, Rehovot, Israel), utilizing the Applied Biosystems DNA sequencer with the ABI BigDye Terminator cycle sequencing kit (Applied Biosystems, Foster City, CA).

Sequence editing, consensus, and alignment construction were performed using Lasergene software, version 5.06/5.51, 2003 (DNAStar, Inc., Madison, WI) and Geneious software version R9 (https://www.geneious.com/academic/). The final nucleotide sequences of the resulting amplicons were compared with data deposited in GenBank. *Mycoplasma* species delineation threshold was 97% as proposed previously(Volokhov *et al*. 2011).

### 2.4. Movement data

To quantify the movement and behavior of Griffons, we deployed GPS and accelerometers tags provided by ’Ornitela’ (Ornitrack 50 3G transmitters). These transmitters, weighing a mere 50 grams (<1% of the Griffon’s body mass), were affixed using a Teflon harness to the lower back of the Griffons in a leg-loop configuration. The GPS tags offer precise spatial positions (typically ±5 m error) and allow for extensive data storage. Notably, these solar-powered tags communicate data via cellular (GSM) networks, facilitating near-real-time tracking (settings include rate of GSM data communication). This immediate identification feature proves valuable for the INPA rangers, enabling prompt actions when necessary, such as in cases of suspected poisoning when Griffons land in high-risk zones. This technology has already been instrumental in mitigating the severe effects of two poisonings in 2021.

These tags have the capability to provide data for up to 3 years and are programmed to record fix locations every 10 minutes, with reduced sampling effort during periods of low battery power (e.g. every hour in cloudy conditions preventing sufficient solar re-charging).

Between 2020-2022, we GPS-tracked 90 wild-born, free-roaming Griffons in the Negev that had been also sampled for pathogens. Here, we investigate the potential effect of mycoplasma on movement, hence we include only a two-week period (1-14 days after sampling) of their tracking. Two weeks is usually minimum duration of infection by mycoplasma infection, despite considerable variation between animal and human patients and species, from several days to even years (Sydenstricker *et al*. 2006; Mok *et al*. 2008; Nilsson *et al*. 2008; Plowright *et al*. 2017; Feberwee *et al*. 2022). To ensure the reliability of our tracking data we excluded days with less than 40 GPS locations per day, and days the Griffons were inside the capture cage (some individuals are re-trapped during the relevant two weeks period). We then filtered out Griffons with less than two reliable days. For the season of 2019 and for the sites of the Carmel and the Golan, sample sizes were too low and were excluded.

We have used two main indices of flight performance to evaluate mycoplasma’s effect on Griffons. First, a Griffon was considered to be flying if at least two consecutive GPS fixes (with up to 25 min between them) had a ground speed of over 4m/s. We then quantified the likelihood of flying each day using 1 h as the minimal threshold (a logical variable, to indicate flight). Second, we have used total distance flown per day for the subset of ‘flight’ (true) days (sum of the distance between all flight fixes per day, in km). We analyzed how these parameters were dependent upon mycoplasma disease status (positive or negative for mycoplasma by PCR analysis), while accounting also for the other confounding factors as predictors in the model: age, origin, status, sampling site (Carmel vs Negev). Similarly, we also analyzed other parameters describing the daily movement but these will be only briefly discussed in this chapter. Namely, we quantified and modeled the 1) Maximum displacement- the linear distance between the starting point and the most distant point recorded in a day (km); 2) Average and 3) maximal daily flight speed across all ‘flight’ fixes (km/hr); 4) Flight duration- sum of the time spent flying (hrs/day). We also looked on the timing of Griffons flight (using Suncalc package in R environment, Thieurmel & Elmarhraoui 2022), logging the 5) take off and 6) return times (hours)- calculated as the number of hours after sunrise/before sunset for the first/last flight fix, respectively. Finally, in addition to daily indices we also examined flight segments of at least three consecutive fixes and calculated their average segment distance (length in km) and duration (in hrs).

### 2.5. Statistical analysis

All data manipulation and analysis was done with R Software, (R Core Team 2018). We used the glmmTMB package for model analysis (Kristensen *et al*. 2016; Brooks *et al*. 2017), Akaike’s Information Criteria for model selection (Anderson *et al*. 2000), ‘*emmeans*’ for post-hoc comparison (Lenth 2023), and the Dharma package for the residual diagnostics of the models (Hartig 2022). Calculations of movement behaviors used the R packages ‘*geosphere*’ (Hijmans 2022), ‘*raster*’ (Hijmans & van Etten 2012), ‘*move*’ (Kranstauber *et al*. 2011), and ‘*mapproj*’ (McIlroy 2023).

For addressing *H.A1* regarding predictors of infection (i.e., likelihood of being positive for *Mycoplasma* spp., a Boolean outcome variable), we used logistic regression models with the year as a random factor (GLMM- General linear mixed model, Binomial family distribution, with a link-logit function). NMDS, CCA and RDA were not used because they are traditionally used for continuous data, and we could not use them without data transformation. The categorical predictors examined were: 1) sex; 2) age group (juvenile, sub-adult, adult); 3) origin (wild-born, breeding program-born, imported); 4) status (captive breeding-program, captive imported, free-roaming); 5) sample site (Negev, Jerusalem, Carmel, Golan), and their interactions. The sample size and collinearity prevented us from creating a full model with all variables combined (origin, site, and status could not be used together due to their high overlap), so each of these variables was considered separately and (when possible) in additive combinations. Because of the structure of the dataset, we used different subsets for different predictors. First, we tested the effect of origin, status, and sampling sites on the data from 2019 and 2020 because no imports occurred in 2021-2022. Furthermore, since these models indicated age as a central predictor, we used a subset of Negev Griffons (in all the years) to test the effect of this predictor on a homogenous group (same origin, status, and site) and the largest subset (N=198 samples, 127 individuals), thus mitigating possible confounding factors.

Testing predictions regarding our second hypothesis (*H.B1*), we used a chi-square test to differentiate between observed and expected proportions for the *Mycoplasma* species in the different age groups and different origins. We did so for the six most prevalent *Mycoplasma* species in our study: *M.* sp. strain 005v*, M. vulturii, M. synoviae, M.* sp. strain 18b*, M.* sp. strain MV120BA & *M.* sp. strain 47802B (at decreasing order of prevalence).

We then compared *Mycoplasma* spp. diversity among status and age groups using three commonly used indices: richness (number of species), Shannon diversity (H; Eq 1) and Simpson index (D; Eq 2).

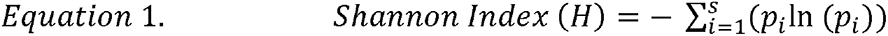

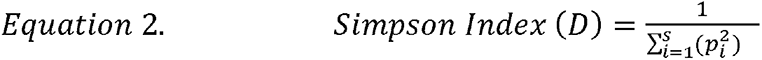

*p* is the proportion of individuals of the *i*-th *Mycoplasma* species found (n) divided by the total number of positive individuals (N), summed (Σ) over *S* species, and *ln* is the natural log. The Simpson index is a dominance index because it gives more weight to common or dominant species. In this case, a few rare species with only a few representatives will not affect the diversity.

Our data has substantial differences in sample size among status groups, with many (119) free-roaming compared to merely 8 breeding program and 14 imported Griffons. To overcome the effect of these differences on *Mycoplasma* species diversity we used a boot-strapping randomization of the wild-born samples for comparing diversity indices (number of species, Shannon index and Simpson index) among groups. Specifically, we determined if the observed indices for the imported and breeding-program Griffons differed significantly (at 95% confidence) from the diversity in a random subset of 8 free-roaming Griffons, permutated 1000 times. Similarly, we compared diversity indices between age groups (29 juveniles, 57 sub-adults and 55 adult Griffons) using bootstrapping of the two latter (larger) groups. We contrasted the observed juveniles’ diversity to the diversity of a random sample of 29 Griffons (either adults or sub-adults, separately), and repeated this procedure for 1000 random permutations to determine if the juveniles significantly differed in the three diversity indices.

The analysis of mycoplasma’s effect on movement was done with a GLMM with a binomial distribution (link-logit function). The Boolean response (flight/no-flight) was a function of mycoplasma infection status (Pos/Neg), as well as age, sex, and their interactions as predictors. Individual identity and year were considered as random factors to account for repeated measures. Similarly, a GLMM with a Gamma distribution (log-link function) and the same structure and predictors was used for comparing total daily distance (a continuous positive explained factor), as well as for the other parameters: maximum displacement, average segment, average segment time, flight time, average flight speed- per day, maximum flight speed, take off time and return time.

## 3. Results

### 3.1. The effect of vulture traits and sampling site on mycoplasma prevalence

Our dataset includes 244 samples from 167 Griffons sampled between 2019-2022. None of the vultures showed clinical signs of disease or were positive for the other pathogens checked (New Castle Disease virus, West Nile virus, Avian Influenza or Chlamydia). The prevalence of *Mycoplasma* spp. for all the samples together was 0.7 (N=170 positives in total) and differed between the years (IZ^2^ = 14.663, df = 3, *p*<0.01, age groups (IZ^2^ = 9.0917, df = 2*, p<* 0.01), and Griffon status (X^2^ = 6.2017, df = 2, *p*<0.05). The prevalence was lower in adults and higher in juveniles (0.61 for adults (N=109 samples), 0.74 for subadults (N=92 samples), and 0.84 for juveniles (N=43 samples); a posthoc analysis confirmed that adults and juveniles were significantly different in prevalence (FDR-adjusted p-value, *p*<0.05). The captive imported Griffons had the highest prevalence (0.95, N=19 samples), while the prevalence in captive breeding program Griffons was lower with 0.64 (N=14 samples), and for free-roaming Griffons was 0.68 (N=73 samples). The prevalence of mycoplasma in captive imported Griffons was significantly different in posthoc analysis from free-roaming (FDR-adjusted p-value, *p*<0.05).

To further understand the effect of each of the predictors on mycoplasma prevalence (age, origin, status and location), and account for possible dependencies among them we supported this simple analysis with generalized linear mixed models, accounting for the years as random effects. The leading model (AICc weight of 48%) included the additive effects of status and age. Age was included also in the following two models as a lone predictor (AICc weight of 34%) or with origin (AICc weight of 11%) (Appendix, **TableA1**). Predicted effect sizes from the leading model agree with the above analysis, showing that juvenile Griffons have higher *Mycoplasma* spp. prevalence than adult Griffons (Juveniles> Subadults>Adults), and captive imported Griffons had a marginally higher prevalence than free-roaming Griffons (**Fig. 2**; Appendix, **TableA2**).

**Figure 2.**
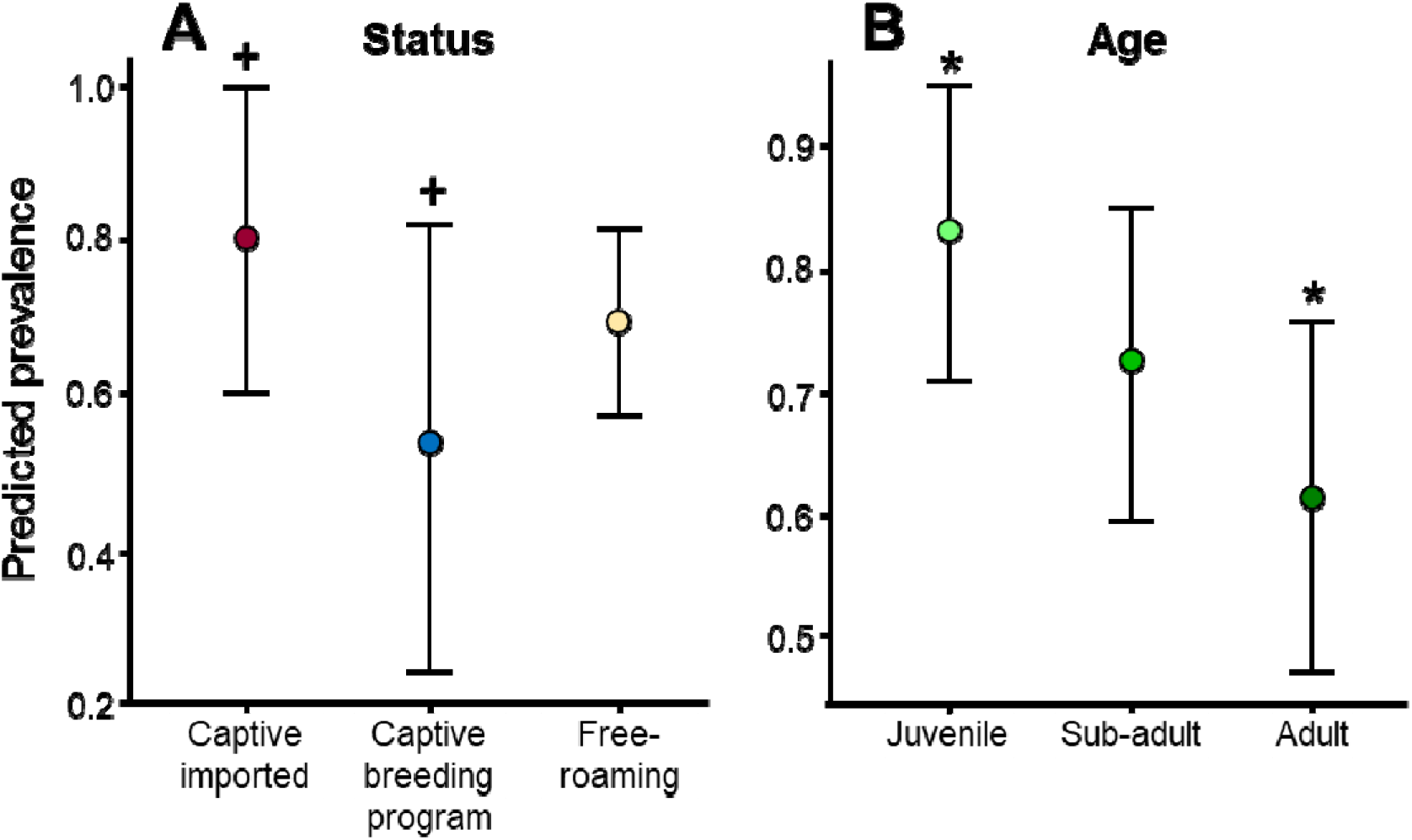
The predicted effects of status (**A**) and age (**B**) on the prevalence of *Mycoplasma* spp. (N = 244 samples from 167 unique individuals, some (N=64) individuals were sampled repeatedly but not in the same year). *P <0.05 for adult compared to juvenile, ^+^P =0.05 marginally significant difference for captive imported Griffons compared to captive breeding program Griffons).

Focusing on a subset of the early years (2019-2020) for evaluating the effects of origin, status and location (N =110 samples, 106 individuals) we found that the leading models predicting the prevalence of *Mycoplasma* spp. included origin (Top model; AICc weight of 24%), age (second model; AICc weight of 23%), status (third model; AICc weight of 19%), and age and status together (fourth model; AICc weight of 11%) (Appendix, **Table A3**). Focusing on the Negev samples subset to examine more accurately the effects of age (N =198 samples, 127 individuals) we found similar results, with the model including age as a predictor having an AICc weight of 87% compared to the null model. Predicted effect sizes, show that juvenile Griffons have higher *Mycoplasma* spp. prevalence than adult Griffons (Juveniles> Subadults>Adults) supporting the analysis of the full data.

### 3.2. Differences in Mycoplasma species distribution and diversity indices between Griffons from different origins or of different ages

Sequence analysis of 137 out of 170 samples positive for *Mycoplasma* spp. revealed nine different species (**Fig. 3**):

1. *Mycoplasma* sp. strain 005v (total N=42).
2. *Mycoplasma vulturii* (total N=28)
3. *Mycoplasma synovia* (total N=26)
4. *Mycoplasma* sp. strain 18b (total N=21)
5. *Mycoplasma* sp. strain MV120BA (total N=9)
6. *Mycoplasma* sp. strain 47802B (total N=8, all samples were with equal homology to *Mycoplasma* sp. strain *M222-5*)
7. *Mycoplasma arginini* (total N=1)
8. *Mycoplasma gypsis* (total N=1)
9. *Mycoplasma maculosum* (total N=1)

**Figure 3.**
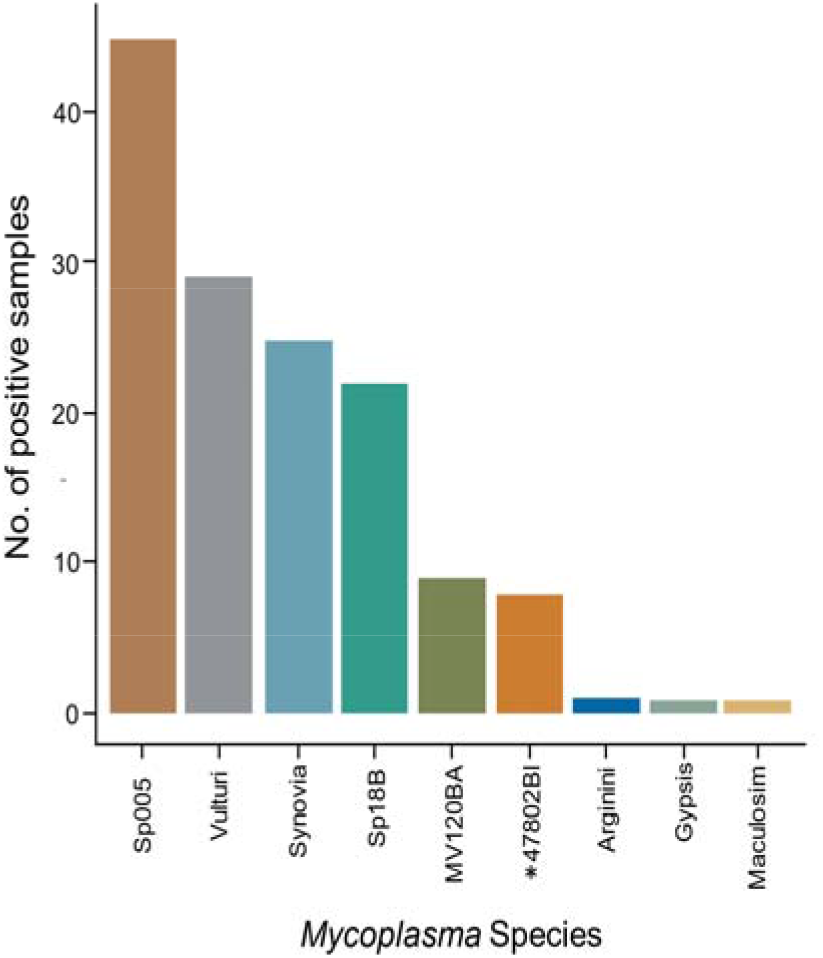
Bar plot of the number of samples found for each species (N=137 samples). Colors represent the different species. \**M.* sp. 47802B or M222-5

(The other 33 positive samples either showed low homology to any species known, had multiple peaks in the sequence or did not have enough genetic material) Using a subset of the six more common species of *Mycoplasma* (*M.* sp. 005, *M. vulturii, M. synoviae, M.* sp. 18b, *M.* sp. MV120BA & *M.* sp. 47802B), we found significant difference species compositions for the different Origins, Status of Griffons, Sampling location and Age, while no significant difference was identified for the different years. Juveniles differed significantly from the adults (Difference between age groups, X^2^= 26.394, df = 10, *p*< 0.01, FDR-adjusted p-value for post hoc between adults and juveniles P<0.001). Interestingly, one species increased with age *M. sp. 005,* whereas another species *M. vulturii,* decreased with age (**Fig. 4A**). We also found a significant difference in species composition between the different status groups of Griffons (X^2^ = 63.21, df = 10, P<0.001). Species composition also differed significantly in captive Imported Griffons from both captive breeding program Griffons and free-roaming Griffons (FDR-adjusted p-value P<0.01, and P<0.001, respectively), and between the two latter groups themselves (FDR-adjusted p-value P<0.05).

**Figure 4.**
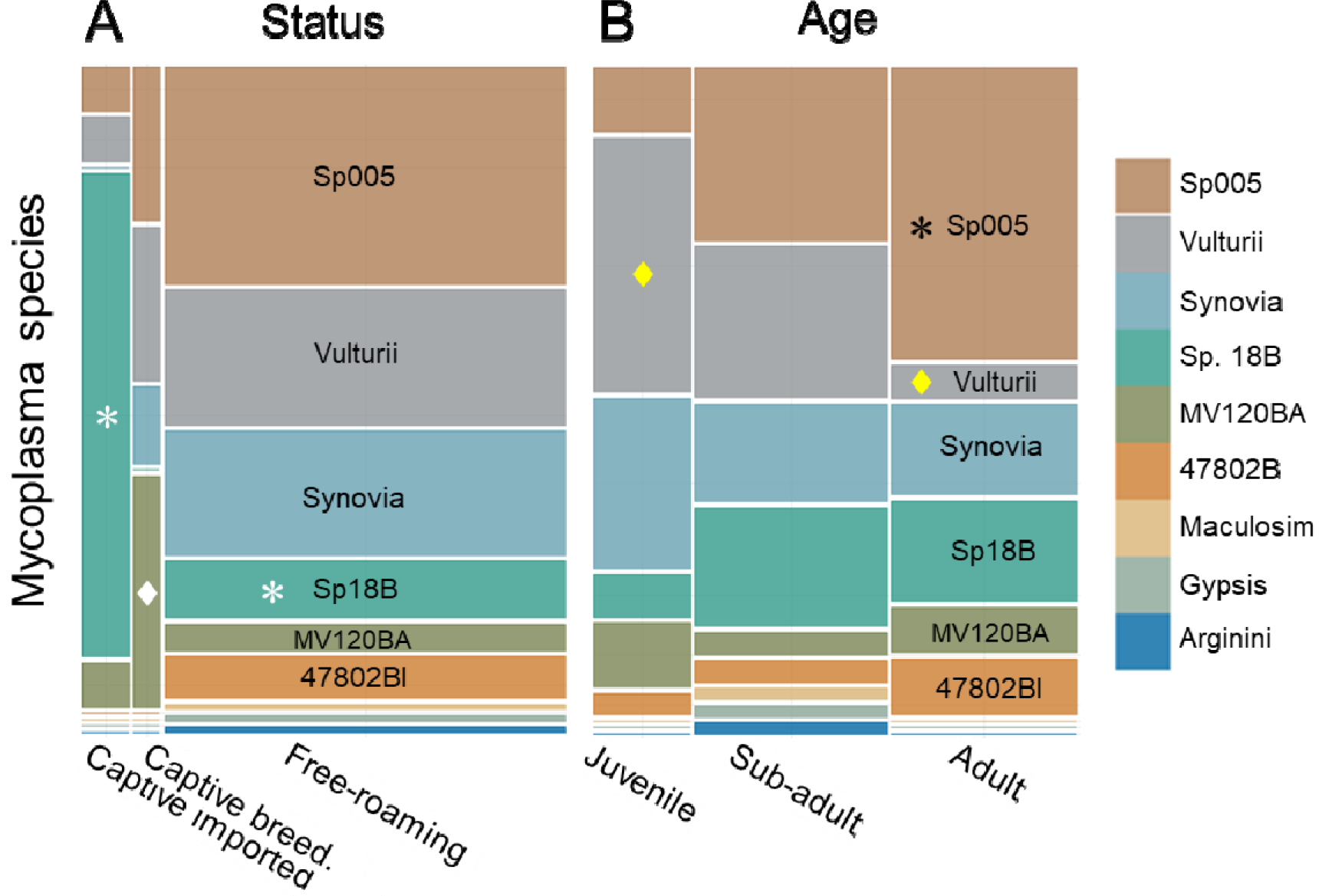
Mosaic plot of the *Mycoplasma* species composition. Each color represents a different species. **A.** Species composition for the different statuses. The width of the x-axis represents the group’s sample size (imported N=14, breeding-program born N=8, wild-born N=119). The species (color) heights (the y-axis) represent their proportion from the group samples. The white diamond in *M.* sp. strain MV120BA represents significant difference between captive breeding program Griffons compared to both captive- imported and free-roaming Griffons. The white asterix in *M.* sp. strain 18b species represents its significantly high abundance in the captive imported Griffons compared to wild-born Griffons **B.** Species composition for the different age groups. The width of the x-axis represents the number of samples for each group (juveniles N=29, subadults N=57, adults N=55). The length of the different species (color) in the y-axis represents the proportion of this species out of the samples for each age group. The yellow diamond in the species *M. vulturii* represents the decrease in the abundance of this species with age. The black asterix in *M.* sp. strain 005 represents the increase in the abundance of this species with age.

Specifically, the *M.* MV120BA found to be more common in the captive breeding program Griffons compared to the (captive) imported and the free-roaming Griffons. In contrast, both *M.* sp. 005 and *M. vulturii* were more common in the free-roaming and the breeding program-born Griffons, whereas *M.* sp. strain 18b was much more common in the Griffons imported from Spain (**Fig. 4B**). Indeed, the majority (11 out of 14 (79%)) of the sequenced samples for imported vultures were identified as this species(18b): six of these were sampled in the quarantine in Jerusalem zoo just after import in 2019, and another five were sampled in captivity in the Carmel and Golan in 2019-2020. Only one breeding-program born Griffon was found positive with this strain (Carmel on 2019). This strain was only rarely found in the wild-born Griffons (9 out of 118, 7.6%): with none in wild on 2019, one from 2020 and eight from 2021-2022, presumably after it was disseminated into the population by released captive vultures.

The diversity of species differed between the different status groups (Appendix, **Table A4**) and age groups (Appendix, **Table A5**). The captive imported group (N=14) had a low diversity with richness of merely four *Mycoplasma* species, H=0.755 and D=0.367. These values were lower than the observed for the free-roaming Griffons (N=119), with richness of nine species, H=1.678 and D=0.772. The bootstrapping showed that the latter group remains significantly higher, with diversity indices of H= 1.286 ± 0.24 (P<0.05 and D=0.683 ± 0.087 P<0.05) even while accounting for a smaller sample size. Griffons in the captive breeding program had intermediate diversity with richness of four species, H=1.321 D=0.719, that did not differ significantly from either group (**Fig 5** **A, B**).

**Figure 5.**
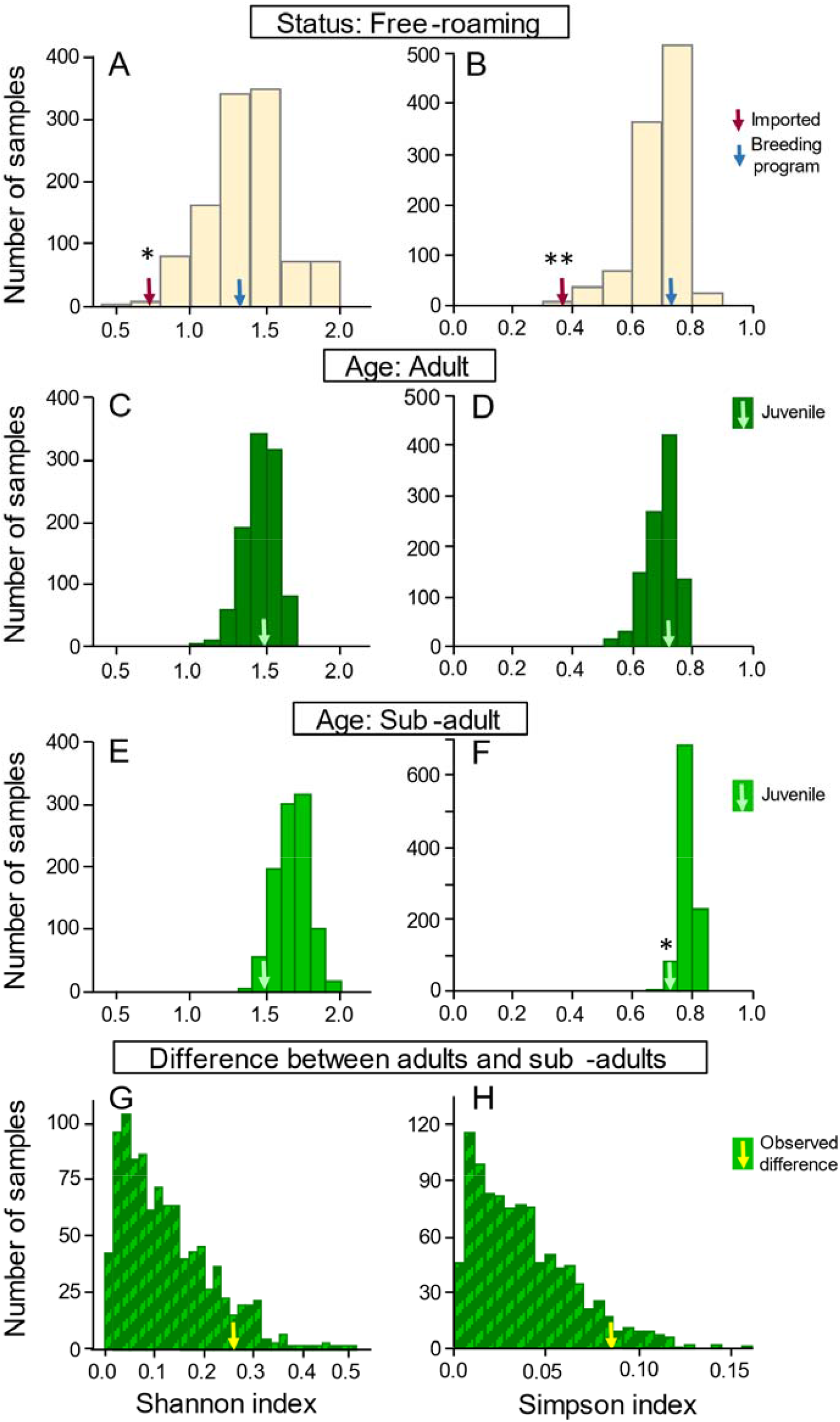
Boot strap randomization to compare *Mycoplasma* species diversity indices (left panels showing Shannon’s H: A, C, E and G; right panels showing Simpson’s D: B, D, F and H) among groups of different status (A and B) and Ages (C-H). Each panel shows a histogram for an index calculated from 1000 random subsamples. **(A & B)** subsamples of 8 Griffons from the free-roaming group show a significantly higher value compared to the observed diversity of the imported Griffons (red arrow) and no difference from the breeding program (blue arrow). **(C & D)** subsamples of 29 adult Griffons show no difference compared to the observed diversity of the juvenile group (N=29, green arrows). **(E & F)** a similar comparison of 29 subsampled sub-adults with the juvenile group shows a marginally significant difference in the Shannon index and a significant difference in the Simpson index. **(G & H)** Histogram of the differences between adults and sub-adults contrasted with the observed differences among these two groups, after a subsampling correction. Observed differences show a trend of higher diversity in the subadult groups but these were not significant for the Shannon index and only marginally significant for the Simpson one.

The diversity of species also differed between the different age groups. Sub-adults had higher diversity (N=57, richness= 9, H=1.758 and D=0.795) compared to juveniles (N=29, richness=6, H=1.499 and D=0.725) and Adults (N=55, richness= 6, H=1.498 and D=0.712). The randomization shows that the difference between subsampled sub-adults (N=29, richness=7) and the juveniles was not significant for Shannon (1.670 ± 0.117; *P*=0.07) but was significant for Simpson (0.781 ± 0.023; P <0.01). Comparing sub-adults to adults also shows that the observed differences are not significant (for observed difference H=0.26 for randomization H=0.123 ± 0.095; P=0.098; for observed D=0.083, for randomization 0.036 ± 0.028; P=0.073). Finally, adults and juveniles had similar diversity indices once corrected for differences in sample sizes through the randomization (N=29, richness= 6, H=1.447 ± 0.108 and D=0.698 ± 0.046).

### 3.3. Does infection (e.g., mycoplasma) affect the movement of Griffons?

We tracked 90 wild-born free-roaming Griffons in the Negev for up to two weeks after their *mycoplasma* sampling, resulting in 116 “tracking periods”. These included 17 juveniles (17 sample periods), each of 10.9 ± 2.74 Griffon days (Mean ± SD); and a total of 185 Griffon days; 31 subadults (39 tracking periods) of 10.4 ± 3.94 days and a total of 405 Griffon days; and 49 adults (60 sampling periods) for 11.4 ± 2.72 Griffon days and a total of 682 Griffon days.

We found that age, sex, and *Mycoplasma* spp. infection affected Griffon’s movement behavior. The best model (AICc weight of 0.23) for the daily probability of flying included age, but not sex or infection. Age was also included in the following two models with the sex of the Griffons (Second best model; AICc weight of 12%) and with infected vs. non-infected (third ranked model; AICc weight of 11%) (Appendix , **Table A4**).

In agreement with our third (alternative) hypothesis (*H.C2*) infection did affect movement distance and duration. For the total distance of flying per day (excluding ‘no-flight’ days), the best model (AICc weight of 0.92) included the interaction between the age and mycoplasma infection status. We found that juvenile Griffons infected with mycoplasma flew significantly lower distances per day than uninfected sub-adults (z. ratio= 5.076, *p*<0.0001), uninfected adults (z. ratio= 3.113, *p*<0.05) and infected adults (z. ratio= -3.261, *p*<0.05) (**Fig. 6A**). Infected sub- adults flew significantly lower distances per day than uninfected sub-adults (z. ratio= 2.986, *p*<0.05). There was no difference between uninfected and infected adults. Similarly, we also found that the best model (AICc weight of 0.81) for flight duration (total hours of flying per day, excluding ‘no-flight’ days), was the interaction between age and mycoplasma infection status.

**Figure 6.**
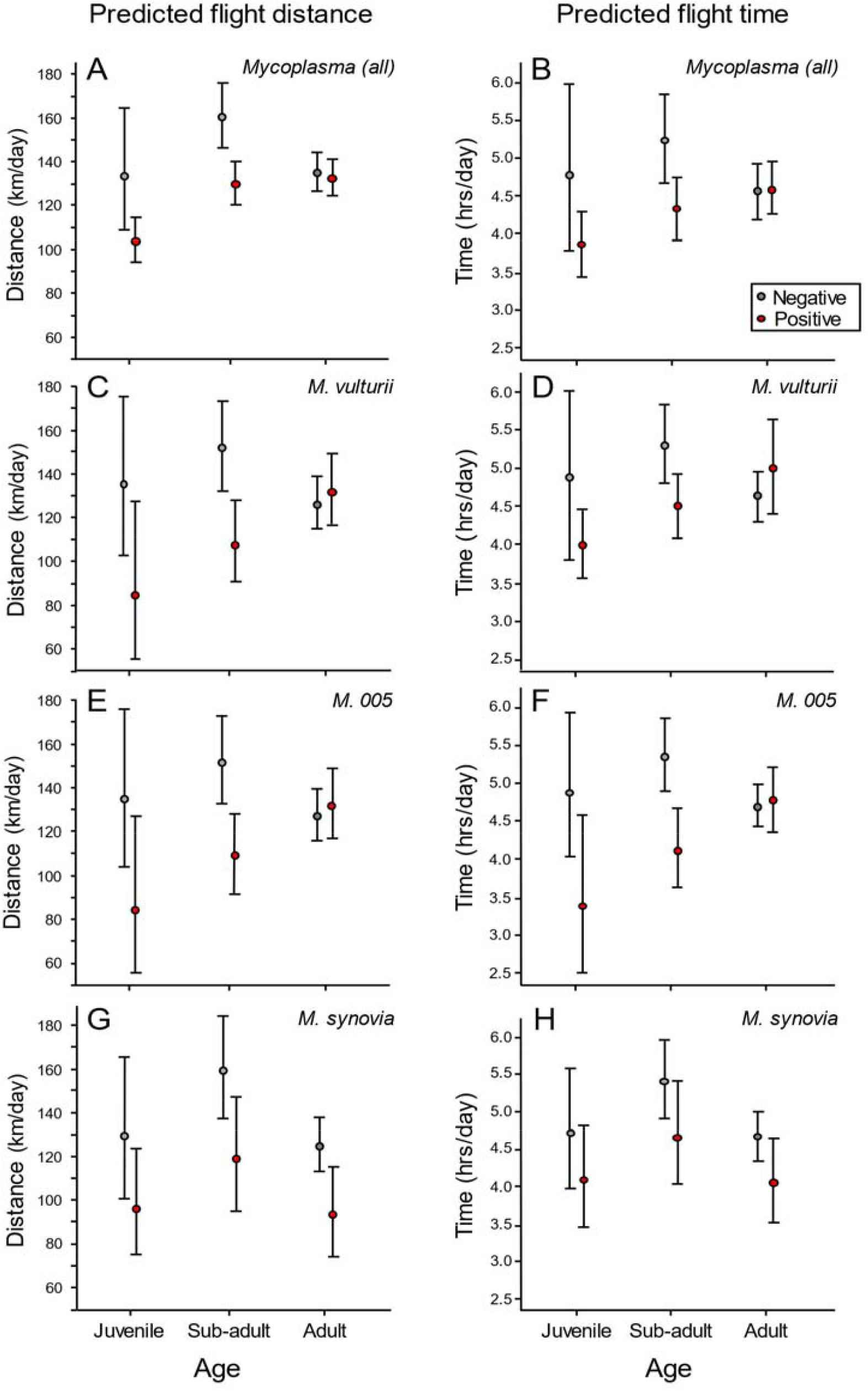
The effect of age and infection on the flight distance (left, panels **A**, **C**, **E** and **G**) and duration (right, panels **B**, **D**, **F** and **H**) of free roaming Griffons. Red and grey dots represent model predicted effects (±SD) for mycoplasma positive and negative groups, respectively. Upper row (**A** and **B**) shows results for all *Mycoplasma* spp. together, while subsequent rows show species specific results for the three most common species, with *M. vulturii* (**C** and **D**), *M.* 005 (**E** and **F**) and *M. synovia* (**G** and **H**). Overall, results show an interactive effect of age and infection on Griffon daily flights, with adults flying less than juveniles and sub-adults, and positive individuals flying less than negative ones. Juveniles and sub-adults show stronger effects of infection, while adults show largely no difference by infection. Similar results are found for the two common species (*M. vulturii, M. 005*), whereas for *M. synovia* infection has non interactive effect, with lower movement across all age groups.

Infected Juvenile and sub-adult Griffons flew significantly less hours per day than **uninfected** sub-adults (z. ratio= 4.586, *p*<0.0001, z. ratio= -2.833, *p*=0.05, respectively, **Fig. 6B**), and marginally different from infected adults. There was no difference between uninfected and infected adults.

We found similar results when we compared infection for the top three *Mycoplasma* species: *M. vulturii, M.* 005 and *M. synoviae* (overall, 27,21, and 11 positive cases, respectively while excluding Griffons positive for alternative species). For both *M. vulturii* or *M.* 005, the best model for the total distance of flying per day and flight duration included the interaction between the age of the Griffons, and mycoplasma status*. M. vulturii*: Juveniles infected with this species flew significantly lower distances per day than uninfected sub-adults (z. ratio= 4.415, *p*<0.001) and infected adults (z. ratio= -3.168, *p*<0.05). Infected sub-adults flew significantly lower distances per day than uninfected sub-adults (z. ratio= 3.921, *p*<0.01), and infected adults (z. ratio= -3.026, *p*<0.05) (**Fig. 6C**). Juveniles and sub-adults infected with this species also flew less hrs. compared to non-infected sub-adults (z. ratio= 4.233, *p*<0.001, z. ratio= 3.399, *p*<0.01, respectively) (**Fig. 6D**). *M. 005*: Juveniles and sub-adults infected with *M. 005* also flew less distance ad fewer hours compared to uninfected sub-adults (**Figs. 6E** **& 6F**). Finally, in contrast to the latter two, for *M. synovia*e the best model included age and infection status (without interaction), possibly reflecting the smaller sample size and limited statistical power. Here, Griffons infected with this species flew significantly less than non-infected Griffons, regardless the age group (z. ratio= 2.478, *p*<0.05) (**Fig. 6G** **& 6H**).

Other flight parameters (maximum displacement, average daily speed and return hour) were also affected by the age of Griffons in interaction with mycoplasma status. Maximum displacement was lower in infected juveniles and sub-adults, the average speed was lower and they returned earlier to the roost site (Appendix, **Fig.A1-3**).

## 4. Discussion

In this study we found high prevalence of *Mycoplasma* spp. (70%) in the Griffon vulture population. We detected high genetic similarities (>95%) to nine different species, six of which have not been previously found in vultures, to my knowledge. This high prevalence and rich diversity, along with measurable effects on Griffons’ behavior highlight the potential relevance of this agent for their ecology and management. Age was correlated with mycoplasma prevalence; as we predicted, juveniles had a significantly higher prevalence than adults, emphasizing the higher sensitivity of young animals to mycoplasma infections. We also found that different age groups differed significantly in their species composition and diversity.

Importantly, our findings support our hypothesis that imported Griffons had a higher prevalence of mycoplasma than local free-roaming Griffons and local breeding program-born Griffons in captivity. Furthermore, imported Griffons differed in species composition and diversity from the local Griffons, and time difference in species appearance may imply that some of the species were introduced into the population after the import. These results emphasize the importance of sampling for parasites and pathogens (here *Mycoplasma* spp. specifically) when transferring animals from region to region and from captive breeding to the wild.

Another support for the ecological importance of mycoplasma infection is the apparent effect on their movement. Our analysis shows that young Griffons (juveniles and sub-adults) infected with *Mycoplasma* spp., flew shorter distances and less time per day, and the results were consistent across the most abundant species *M. vulturii* and *M. 005.* Griffons infected with *M. synoviae* also had reduced flight distance and time per day. These results show that mycoplasma infection influences Griffon movement, even though other clinical signs are not observed, or are hard to identify in routine monitoring.

### 4.1. Mycoplasma prevalence in Griffons

Mycoplasma prevalence in our study was found to be much higher (70%) than in previous research on Griffon vultures in Israel, where 47% and 46% prevalence was identified in Griffons in the wild in Griffons in captivity, respectively (Blass *et al*. 2012). This can reflect a better detection methodology (different primers used), our larger sample size and better sensitivity, or a real change in abundance caused by the reintroduction or by other factors. In free-roaming raptors in Germany, a mycoplasma prevalence of 91% in nestlings and 94% in adult birds was detected (Lierz *et al*. 2008a). However, mycoplasma prevalence in raptors varied according to the species and age groups (Poveda *et al*. 1990a; Lierz *et al*. 2000b, 2002, 2008a).

### 4.2. Mycoplasma species found

Based on the partial sequencing of the 16S-rRNA, nine species were found in the Griffons with highest identity to *M.* sp. strain 005v*, M. vulturii, M. synoviae, M.* sp. strain 18b, *M.* sp. strain MV120BA*, M.* sp. strain 47802B/M222-5, *M. arginini*, *M. gypsis* and *M. maculosum*. Six of these species have not been previously detected in Griffon vultures in Israel (Blass *et al*. 2012) or other regions (Sumithra *et al*. 2013; Plaza *et al*. 2020). Some *Mycoplasma* species are known to cause disease in vultures including respiratory disease (Poveda *et al*. 1990b; Oaks *et al*. 2004; Loria *et al*. 2008; Lecis *et al*. 2010), as well as orthopedic disease (Panangala *et al*. 1993; Ruder *et al*. 2009), however the effect of most species still remains unknown (Lierz *et al*. 2002, 2008a; Lecis *et al*. 2011).

In our research, 42 out of 137 of the sequenced samples had 99-100% identity per sequence compared to *M.* sp. strain 005V, (GenBank accession MT721031). To the best of our knowledge, this species, was originally isolated from a Spanish booted eagle’s trachea (*Hieraaetus pennatus*) in 2005, and has been also isolated from a Cinereous vulture (*Aegypius monachus*), Bonelli’s eagle *Aquila fasciata*, and Short-toed snake eagle (*Circaetus gallicus*) (Joachim Spergser, personal communication). 14 of the sequenced samples also had high homology to *M.* sp. nov type strain *1449* (GenBank accession MK561040).

The second most common species was *M. vulturii* (N=29/137), first isolated from the tissues of an Oriental white-backed vulture (*Gyps bengalensis*) found dead from NSAIDs in Pakistan (Oaks *et al*. 2004), *M. synoviae* was the third most common species (25/137) identified., *M. synoviae* is a common pathogen of poultry and has also been found in wildlife (Luttrell *et al*. 1992; Kleven & Ferguson-Noel 2008; Sumithra *et al*. 2013). It may become systemic and cause bronchitis, air sac infection and infectious synovitis (joint infection) (Kleven & Ferguson-Noel 2008). After introduction of vaccination (2018), based on used of live *M. synoviae* H strain, the prevalence of *M. synoviae* is drastically decreased in poultry broilers and breeders in Israel, but its prevalence in the layers is still high.

*M.* sp. 18b (N=22/137) was the fourth most common species detected. This species was first isolated from a white stork’s liver (*Ciconia ciconia*) in Spain (GenBank accession MT721015), but We do not know of the pathogenicity of this strain nor in which other species it may occur; it has not been previously found in vultures.

*M.* sp. strain MV120BA was found in nine vultures. This species was first described in a white-tailed eagle (*Haliaeetus albicilla*) from Germany (GenBank accession OK095372), but have not been previously found in vultures.

The sequence of eight samples demonstrated equal identity to two *Mycoplasma* species*: M.* sp. 47802B (GenBank accession MT721011), previously identified in a common kestrel *(Falco tinnunculus)* from Germany and *M.* sp. M222-5 (GenBank accession EU684058), identified in a lesser kestrel *(Falco naumanni)* from Spain. These species have not been previously been isolated from Accipitridae, but from Falconidae. The PCR product used for species identification in this study is based on the partial sequence of 16S rRNA, which is in this specific region demonstrated 100% identity between *M.* sp. strain 47802B and *M.* sp. strain M222-5. The sequencing *rpoB* gene as well as intergenic 16S rRNA region may help to distinguish between these species.

*M.gypsis, M. arginini,* and *M.maculosum* were all found only once each of our samples. *M. gypis* was first described in Griffon vultures in 1994 (Poveda *et al*. 1994), and has been commonly found in vultures and other birds of prey (Poveda *et al*. 1994; Lierz *et al*. 2008a; Blass *et al*. 2012). Surprisingly, we only had one positive sample for this species in this study. One of the possible explanations of this finding is identification of two mismatches between sequence of MGSO primer and sequence of 16S rRNA of *M. gypis* and that may impact the annealing of the primer and the efficacy of the PCR reaction. *M. arginini* is frequently isolated from healthy and diseased sheep and goats (Deeney *et al*. 2021), found in wild felids (Sumithra *et al*. 2013). Its zoonotic potential was also raised (Prayson *et al*. 2008; Watanabe *et al*. 2012). The *M. arginini* detected in our sampling is probably due to contamination of the swab with sheep or goat remains in the mouth of the Griffon. *M. maculosum* has been commonly found in dogs and has been known to cause fertility and respiratory infections in dogs (Armstrong *et al*. 1970; Tamiozzo 2022) and can also become zoonotic (Heilmann *et al*. 2001). We believe that this positive sample was either a species close to *M. maculosum* or a contamination of the sample.

### 4.3. Age differences in mycoplasma prevalence and species diversity

When analyzing the full dataset, we found a higher prevalence of *Mycoplasma* spp. in juveniles (Griffons on their fledging year, 84%, N=43), compared to sub-adults (1-4 years, 74%, N=92) and adults (> 4 years, 61%, N=109). We found similar results when analyzing a more homogenous subset of Negev free-roaming wild-born Griffons. Lierz *et al*. 2000b had similar to findings of higher prevalence of mycoplasma infection in juvenile raptors. In juveniles of other species, including humans, dogs, ruminants, song birds, poultry and wildlife (Bradbury *et al*. 1988; Altizer *et al*. 2004; Chalker *et al*. 2004; Gabinaitiene *et al*. 2011; Sumithra *et al*. 2013; Song *et al*. 2023), prevalence of *Mycoplasma* spp. also tends to be higher, possibly due to the immaturity of the immune system or vertical transmission (Atkinson & Waites 2014).

Species composition and diversity were found to be different between the different age groups of the Griffons, similar to prevalence. Interestingly, *M.* sp. strain 005v prevalence increased with age, whereas *M. vulturii,* prevalence decreased with age. The sub-adult Griffons showed higher diversity compared to juveniles (significantly) and adults (marginally significant). The prevalence of many *Mycoplasma* species is higher in young animals (Bradbury *et al*. 1988; Gabinaitiene *et al*. 2011; Sumithra *et al*. 2013; Song *et al*. 2023). On the other hand, some species may cause chronic infections and may be detected for many years in the same individual (Haier *et al*. 1999; Dallo & Baseman 2000; Nijs *et al*. 2002), which may be the case for *M. sp.* strain 005v in our research. Increase in species diversity in sub-adults, may be due to most of the imported vultures being from this age group, thus adding the imported species to the local species number. Another possibility is that the sub-adult Griffons travel larger distances (Arkumarev *et al*. 2021) and have more social interactions compared to adults, thus increasing the possibilities for infectious encounters (Acacio *et al. in prep*).

### 4.4. Mycoplasma prevalence and diversity differ between imported and local Griffons

Notably, we found that imported Griffons had higher mycoplasma prevalence compared to both captive breeding-program born Griffons and free-roaming Griffons. When we analyzed a subset for the years 2019-2020, we found that the leading model included origin, *i.e.,* where the Griffon was born: either imported from Catalonia (Spain), captive-born in Israel, or wild-born in Israel.

We found that captive imported Griffons had lower *Mycoplasma* species diversity compared to free-roaming Griffons, despite having a higher prevalence. This might be due to close contact with the same few individuals. The species composition of the imported Griffons differed from both the breeding-program born and free-roaming Griffons. Notably, *M.* sp. strain 18b was much more abundant in the imported Griffons compared to free-roaming Griffons, and only appeared in the latter group later on. *M.* sp. strain 18b was the most common *Mycoplasma* species found in the samples from Griffons imported from Spain (11/14). Eight of these samples were collected a few days after Griffon’s arrival, while in quarantine in the Jerusalem Zoo in 2019, another three were in captivity in the Carmel and Golan in 2019-2020. In the Negev free-roaming Griffons the first positive sample for *M.* sp. strain 18b was in September 2020 and eight more Griffons *M.* sp. strain 18b positive samples were detected in free-roaming Griffons in 2021-2022. These findings might suggest that this strain has been imported from Spain and has spilled over into the Israeli wild Griffon population.

The captive breeding program Griffons did not differ from the free-roaming Griffons in species diversity, however they differed from both the imported Griffons and the free-roaming Griffons in species composition. The most abundant species in this group was *M.* sp. strain MV120BA.

These findings highlight the importance of diagnosis of mycoplasmas when translocating animals, both between different areas and from captivity (i.e., breeding programs and rehabilitation facilities) to the wild. *Mycoplasma* infections are of concern in captive-bred wildlife and have been found in several species, including mammals, reptiles and birds (Hill 1975; Clippinger *et al*. 2000; Friend 2006; Grazziotin *et al*. 2011; Lecis *et al*. 2011; Sumithra *et al*. 2013). The close proximity of individuals during transfer or captivity may increase infection rate, while the stress and acclimation to the new environment may suppress immune activity (Kock *et al*. 2010, 2018; Sainsbury & Vaughan-Higgins 2012) results in high transmission rate. In some cases, latent disease may reemerge due to stress, as has been seen in Cape buffaloes *(Syncerus caffer)* with foot and mouth disease, which infected cattle in the relocation site (Hedger & Condy 1985). Translocations (including reintroductions) have been known to cause spillovers into naïve native populations and domesticated species (Viggers *et al*. 1993; Kock *et al*. 2010). As seen in reintroduced desert tortoises with upper respiratory infection which may have infected the free-roaming population (Jacobson *et al*. 1991, 2014; Brown *et al*. 2004a), the spread of *Plasmodium* spp. to extant wild turkeys (Castle & Christensen 1990), and the spread of rabies in racoons from the southeast to mid-Atlantic US (Jenkins *et al*. 1988). On the other hand, translocated animals are naïve to the local pathogens, and may be more suspectable to disease due to the stress off translocation, as reported in the devastating effect of canine distemper to reintroduced the black-footed ferret (*Mustela nigripes*) in the US (Williams *et al*. 1988).

### 4.5. The effect of mycoplasma infection on the movement behavior of Griffon vultures

Age is known to affect Griffon behavior, with adults largely traveling less then juveniles and sub- adults (Spiegel *et al*. 2015, Acacio in press; Arkumarev *et al*. 2021). By accounting for age and focusing on the two-week period following the mycoplasma sampling we were able to demonstrate that infection results in a reduction of daily flight and duration, and that the effect is age dependent. While infected sub-adults and juveniles flew less (shorter distances and duration) compared to their non-infected counterpart, adults were apparently not affected. This was also found when we examined the effect of each of the most common species of *Mycoplasma* on movement. This result suggests that infection with mycoplasma may influence the health and behavior of Griffons, in contrast to previous research on mycoplasma infection in raptors, that has suggested that most of these species are commensal (Lierz *et al*. 2002, 2007, 2008a). Similar results were found in desert bighorn sheep (*Ovis canadensis nelsoni*), infected with *Mycoplasma ovipneumoniae,* but did not demonstrate clinical signs; infected sheep showed a long-lasting decrease in mean daily movement (Dekelaita *et al*. 2023). Notably, even if the isolation of mycoplasma is from individuals without clinical signs (at least to the level one can find in the relatively superficial examination during captures), it does not mean the species is commensal (Taylor-Robinson 1996). Tracking allows us to evaluate the animal remotely, without the stress of capture and handling and enables us to evaluate changes in behavior such as movement more naturally and for longer periods of time, thus obtaining a better understanding of diseases effects on animals.

While establishing the ecological relevance and impact of *Mycoplasma* spp. our study cannot directly address the consequences, or the mechanism of infection. The differences in flight parameters may also be correlated with reduced fitness, such as lethargy, reduced respiratory ability and arthritis which may occur in pathogenic avian mycoplasma infections (Panangala *et al*. 1993; Erdélyi *et al*. 1999; Ruder *et al*. 2009; Sumithra *et al*. 2013). Presumably focusing on a longer tracking period, rather than the short-term infection period can reveal if the reduction in movement is associated with long term effects. Further, while we focus on movement *following* the sampling in the context of mycoplasma effect on behavior, we cannot prove causality or exclude the alternative explanation that reduced movement (and more time at the communal roost) enhances the probability of infection, and leads to the apparent negative association between movement and infection inferred here as ‘sickness behavior’ (Dougherty *et al*. 2018; Spiegel *et al*. 2022). Similarly, mycoplasma may act as an opportunistic infection with other infections or in cases of poor health and weakened immune system, providing another alternative explanation for our findings (Song *et al*. 2015; Canter *et al*. 2019; Rangroo *et al*. 2022).

### 4.6. Study limitations and future directions

First, we note that *Mycoplasma* species can cause acute or chronic infections. Because most of the individuals were sampled only once a year it was hard to assess the persistence of infection. In the future, it would be interesting to sample the same individuals repeatedly in captivity and in the wild, to assess different *Mycoplasma* species’ persistence and difference in immunity and other health indicators in parallel. Second, our sample size for juveniles was lower than for the other age groups limiting our ability to represent the movement of this group or its different from sub-adults, as these were only juveniles under age of one year. Unfortunately, the population of Griffons in Israel is small, with low recruitment, therefore it would be hard to enlarge this sample size. Third, isolation of mycoplasmas from vultures could significantly upgrade this research. Unfortunately, and despite using multiple protocols, the isolation of these species did not succeed. Future research is needed to improve the identification of the species in the lab. Some of the sequences we found were with similar homology to two species in the GenBank. Sequencing other regions like *rpoB* gene and intragenic 16S rRNA region might give more conclusive results.

Despite the fact that common pathogens of birds such as West Nile, Avian Influenza and New Castle virus were not identified in this study, we cannot exclude the presence of other pathogens which may impact Griffon’s movement like we found for mycoplasmas. An experimental approach targeting these questions can be very informative, especially if combined with subsequent tracking. For instance, treating Griffons with antibiotics for mycoplasma and checking if this alters the change we see in their movement. It would also be interesting to understand if infection with mycoplasma may also influence other fitness metrics, such as fertility, and hatching and fledging success.

### 4.7. Conservation implications and conclusions

In our research we found very high prevalence of mycoplasma in Griffon vultures, including species not yet described in Israel or in Griffon vultures (*M. sp.* strain 005v, *M. sp.* strain 18b, *M. sp.* strain MV120BA, *M. sp.* strain 47802B/M222-5, *M. arginini* and *M. maculosum*. Outbreaks of mycoplasma have become very common in wildlife species and can threaten conservation efforts, highlighting the importance of effective monitoring of pathogens presence in general and mycoplasma specifically (Fischer *et al*. 1997; Besser *et al*. 2012; Sumithra *et al*. 2013; Highland *et al*. 2018). Finding species that were not previously described in vultures suggests that *Mycoplasma* species are not always strictly host-specific (Sumithra *et al*. 2013; Highland *et al*. 2018).

We have found a higher prevalence and presence of different species in imported Griffons in comparison to the local Griffons, with suggestive evidence of introduction of one species into the native population (*M.* sp. strain 18b, found initially only in imported Griffons from Spain, was found a year later in the wild population). This demonstrates that translocation management (here importing vultures from Spain) may introduce pathogenic species across geographical barriers, with enhance risks for the local Griffons in the breeding program and in the wild (Erdélyi *et al*. 1999; Clippinger *et al*. 2000; Grazziotin *et al*. 2011; Lecis *et al*. 2011; Sumithra *et al*. 2013), and to the translocated individuals themselves (Jacobson *et al*. 1991; Brown *et al*. 2002; Sumithra *et al*. 2013), as well as for spill over into other hosts. Given these results, we highlight the need to properly quarantine new animals before introduction to a breeding facility, screening for mycoplasma and other pathogens, isolate and treat infected animals as well as pre-release screening of captive animals into the wild population.

Finally, similar to other species (Lierz *et al*. 2000b), juvenile Griffons had higher prevalence, different species composition and stronger reduction in movement compared to adults, confirming the susceptibility of this group to mycoplasma infections. Even in the absence of clinical signs, Griffons infected with mycoplasma flew less (shorter distances and periods), especially noticeable in the sub-adult group. In the adult group, the infection did not seem to change movement behavior (except for *M. synoviae* infection). These findings show that there might be an effect of mycoplasma infection on movement and therefore these species may not be not pure commensals, as previously suspected. This also highlights the importance of combining tracking information to disease ecology, as we gain more true insights on the effect of disease on animal behavior.

## Acknowledgments

We are grateful to many organizations and people that have devoted their efforts to the conservation of the Griffons, for help with the captures and sampling, and for their comments: The INPA Breeding program staff: S. Simchi, A. Bar-On, L. Lior, E. Maatuf, A. Peretz, E. Zisso and many other rangers and keepers for their sterling role in the conservation of the Griffons. We wish to thank the Israeli wildlife hospital staff and Dr. Roni King of the INPA for their diagnosis and treatment of Griffons. We are indebted to the staff of the Ramat Hanadiv Gardens and Natural Park and the Tisch Zoological garden in Jerusalem (especially N. Avni-Magen, Y. Levy Paz and E. Zisso). We would like to thank A. Uzan, E. Arnon, T. Solomon and many others from The Tel-Aviv University Movement Ecology Laboratory for their assistance in the field and for their valuable advice. We are indebted to A. Berkowitz, A. Lublin, I. Farnushi, S. Mashani from the Division of Avian Diseases, Kimron Veterinary Institute, and to R. Lapid, R. Efrat, E. D’Bastiani, K. Gahm, A. Gancz, G. Kahila-Bargal and Z. Aisenberg for their help and recommendations.

The work was funded by the NSF IOS division 2015662/BSF 2019822 (to OS). NA and GV were supported by a stipend from Yad-Hanadiv. MA was supported by the George S. Wise Postdoctoral Fellowship (Tel Aviv University). None of these funders were involved in any of the phases of the research and manuscript.

