## Supplementary figures for "Factors affecting the individual probability of infection with a prevalent pathogen (*Mycoplasma*) and the effect on Griffon vultures’ movement behavior"

**APPENDIX for Mycoplasma article**

**Table A1. Factors affecting the prevalence of *Mycoplasma* spp. infection in Griffons 2019-2022.** Model rankings are based on Akaike Information Criterion (AICc) for the predictors of Age, Origin, Status, and Sex. Sampling years (2019–2022) is included as a random effect in all models. Only Models with Delta AIC < 4 are presented in this table. It should be noted that all five leading models include age). N=244 samples, 167 individuals.

|  | K | AICc | ΔAICc | AICc | Cum.Wt |
| --- | --- | --- | --- | --- | --- |
| Age + Status | 6 | 290.72 | 0.00 | 0.48 | 0.48 |
| Age | 4 | 291.42 | 0.70 | 0.34 | 0.81 |
| Age + Origin | 6 | 293.71 | 2.99 | 0.11 | 0.92 |

**Table A2. Parameter estimates for the top model used for GLMM describing *Mycoplasma* spp. prevalence** **for all samples 2019-2022** in log: Age + Year (sampling year is a random effect in this model). High effects were found for juveniles (higher prevalence) and imported Griffons (higher prevalence). N=244 samples, 167 individuals.

| Parameter | Estimate | SE | t | p |
| --- | --- | --- | --- | --- |
| Intercept (Imported, Juvenile) | 3.548 | 1.198 | 2.96 | 0.00306 ** |
| Age (Subadult) | -0.889 | 0.549 | -1.62 | 0.105 |
| Age (Adult) | -1.444 | 0.525 | -2.75 | 0.00589** |
| Status (Captive breeding) | -2.361 | 1.228 | -1.92 | 0.05453 |
| Status (Free-roaming) | -1.671 | 1.091 | -1.53 | 0.12569 |

**Table A3. Factors affecting the prevalence of *Mycoplasma* spp. infection in Griffons for 2019-2020.** Model rankings are based on Akaike Information Criterion (AICc) for the predictors of Age, Origin, Status, and Sex. Sampling years (2019–2020) is included as a random effect in all models. Only Models with Delta AIC < 4 are presented in this table. N=110 samples, 106 individuals.

|  | K | AICc | ΔAICc | AICc | Cum.Wt |
| --- | --- | --- | --- | --- | --- |
| Origin | 4 | 127.69 | 0.00 | 0.24 | 0.24 |
| Age | 4 | 127.80 | 0.11 | 0.23 | 0.47 |
| Status | 4 | 128.15 | 0.46 | 0.19 | 0.67 |
| Age + Status | 6 | 129.36 | 1.67 | 0.11 | 0.77 |

**Table A4. *Mycoplasma* species diversity indexes between different status of Griffons**

The number of samples taken from different statuses was not distributed equally, so for the Captive -imported and Captive breeding-program I measured the diversity indices from the samples and compared these to indexes calculated 1000 times using a random sample of 8 free-roaming vultures. For the randomized samples for free-roaming Griffons, number of species is presented as the median, and Shannon and Simpson indices as average± SD.

| **Status** | **Samples (N)** | **No. of Species** | **Shannon Index** | **Simpson Index** |
| --- | --- | --- | --- | --- |
| Captive-Imported  (Observed) | 14 | 4 | 0.755* | 0.367* |
| Captive-Breeding program (Observed) | 8 | 4 | 1.321 | 0.719 |
| Free-Roaming (Observed) | 119 | 9 | 1.678 | 0.772 |
| Free- Roaming (Random) | 8 | 4 (median) | 1.286 ± 0.24* | 0.683 ± 0.087* |

*P<0.05

**Table A5. *Mycoplasma* species diversity indexes between Griffons of different age groups**

The number of samples taken from different age groups was not distributed equally, so for the juveniles I measured the diversity indices from the samples and compared these to indexes calculated 1000 times using a random sample of 29 sub-adult and 29 adult vultures. For the randomized samples number of species is presented as the median, and Shannon and Simpson indices as average ± SD. To compare the sub adult and adult group I compared the calculated difference in the observed Shannon and Simpson indexes to a calculated subtraction between 57 and 55 randomized samples out of the adult and sub-adult combined group.

| **Age-group** | **Samples (N)** | **No. of Species** | **Shannon Index** | **Simpson Index** |
| --- | --- | --- | --- | --- |
| Juvenile (Observed) | 29 | 6 | 1.499 | 0.725* |
| Sub-adult (Observed) | 57 | 9 | 1.758 | 0.795 |
| Sub-adult (Random) | 29 | 7 (median) | 1.670 ± 0.117 | 0.781 ± 0.023* |
| Adult (Observed) | 55 | 6 | 1.498 | 0.712 |
| Adult (Random) | 29 | 6 (median) | 1.447 ± 0.108 | 0.698 ± 0.046 |
| Adult-Sub-adult (Observed) | (55,57) |  | 0.26 | 0.083 |
| Adult-Sub-adult (Random) | (55,57) |  | 0.123 ± 0.095 | 0.036 ± 0.028 |

*P<0.01

**Table A6. Factors affecting the probability of flying per day per Griffon** Model rankings are based on Akaike Information Criterion (AICc) for the predictors of Age, Sex, Status of *Mycoplasma* spp. infection and the interaction between them. Sampling years (2020–2022) and individual id are included as random effects in all models. Only Models with Delta AIC < 4 are presented in this table. N=90 individuals, 116 sampling periods, and a total of 1272 Griffon “days”.

|  | K | AICc | ΔAICc | AICc | Cum.Wt |
| --- | --- | --- | --- | --- | --- |
| Age | 4 | 435.09 | 0.00 | 0.23 | 0.23 |
| Age +Sex | 5 | 436.34 | 1.24 | 0.12 | 0.35 |
| Age + Positive | 5 | 438.39 | 1.43 | 0.11 | 046 |

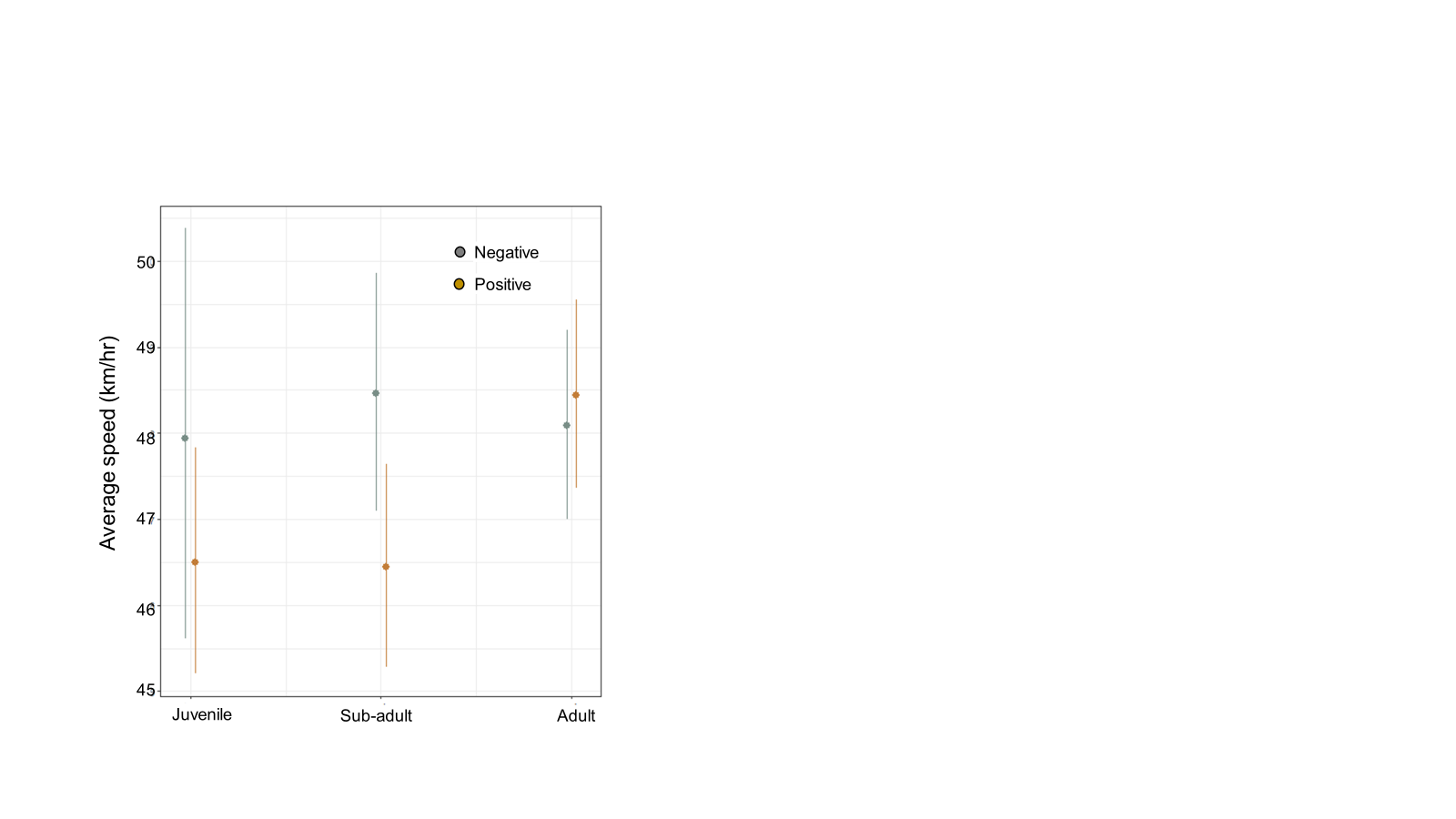

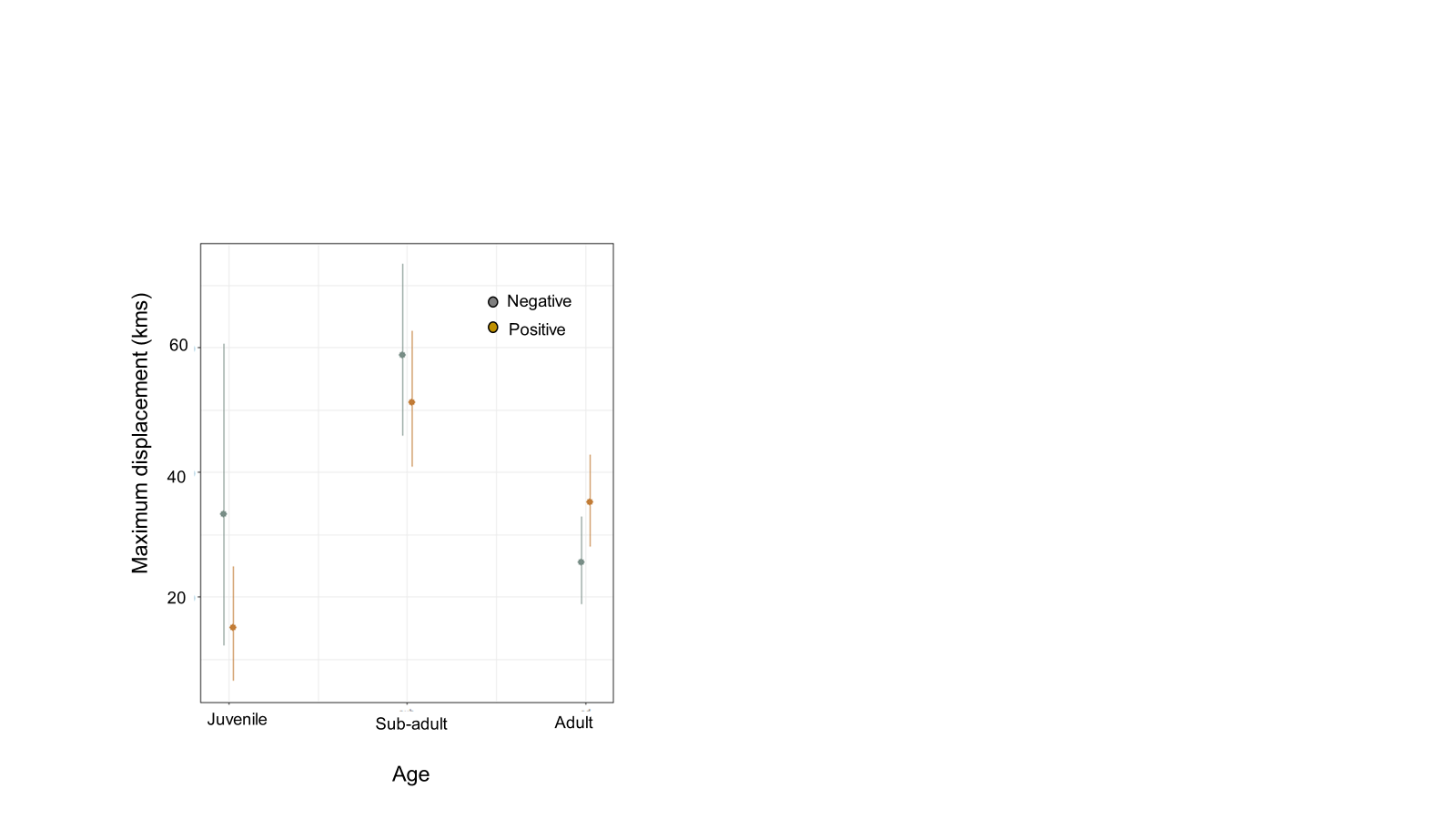
**Figure A1.** The effect of age and infection on the maximum daily displacement (the distance between the beginning point and the furthermost point on the same day). Gold and grey dots represent model predicted effects (±SD) for mycoplasma positive and negative groups, respectively.

**Figure A2.** The effect of age and infection on the average flight speed per day (km/hr). Gold and grey dots represent model predicted effects (±SD) for mycoplasma positive and negative groups, respectively.

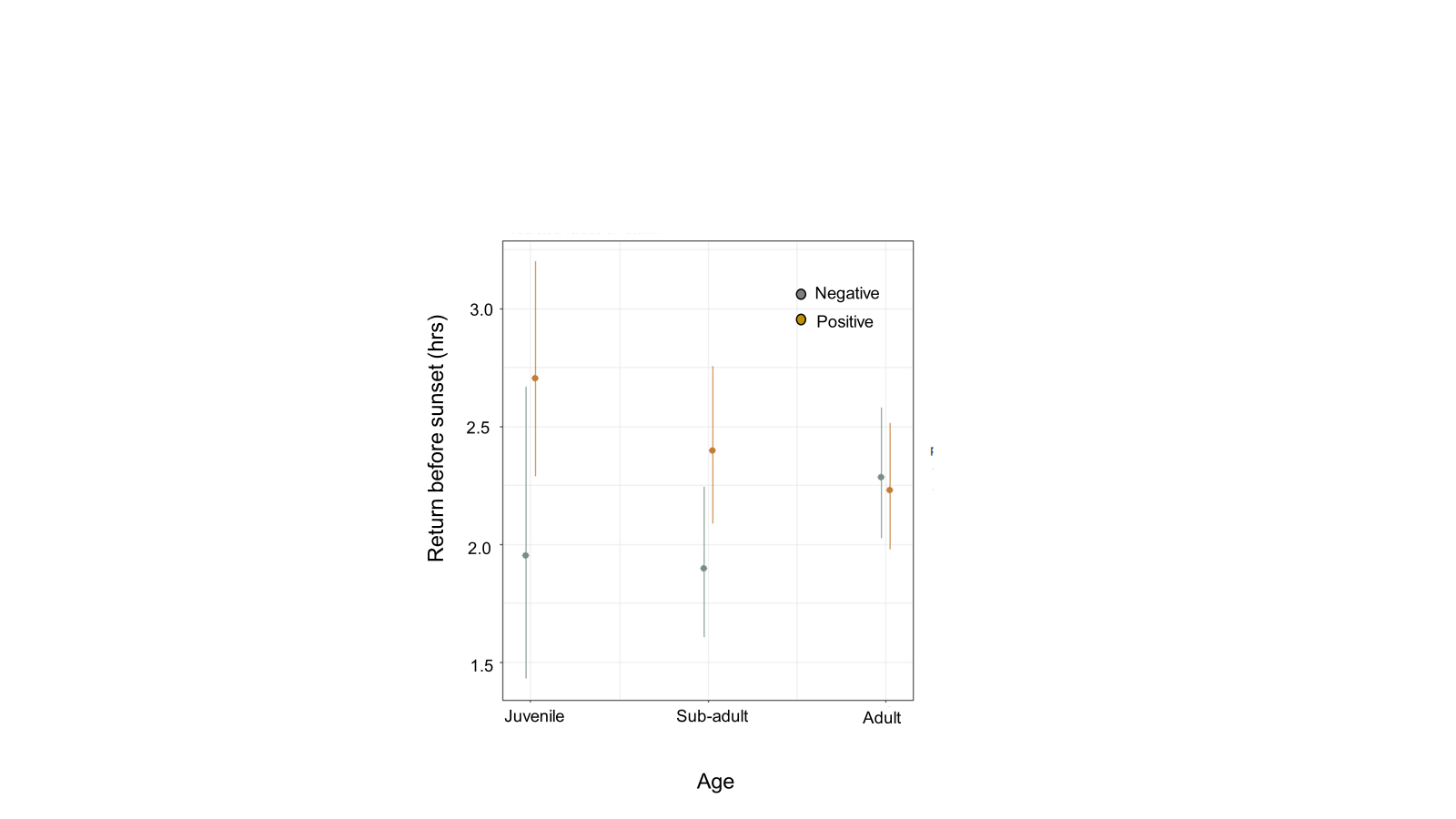

**Figure A3.** The effect of age and infection on the return to roost hour before dawn (end of flight). Gold and grey dots represent model predicted effects (±SD) for mycoplasma positive and negative groups, respectively.
